# Long-read Spatial Transcriptomic Profiling of Patient-derived ccRCC Tumoroids Reveals Heterogeneity in Isoform and Gene Expression

**DOI:** 10.1101/2024.10.16.618643

**Authors:** Mustafa Elshani, Hazem Abdullah, Ying Zhang, Alexander Laird, Peter Mullen, David J. Harrison

**Affiliations:** University of St Andrews, School of Medicine, North Haugh, St Andrews, Fife, KY16 9TF, UK; NuCana plc, 3, Lochside Way, Edinburgh, EH12 9DT, UK; Edinburgh Urological Cancer Group, Institute of Genetics and Molecular Medicine, University of Edinburgh, Western General Hospital, Crewe Road South, Edinburgh EH4 2XU, UK

**Keywords:** spatial transcriptomics, tumoroids, ccRCC, long-reads

## Abstract

Clear cell renal cell carcinoma (ccRCC) is the most prevalent type of kidney cancer characterized by its diverse tumor composition featuring various subclonal populations that hinder effective treatment responses. Tumoroids present an avenue for modelling this diversity and replicating the intricate tumor heterogeneity. Spatial transcriptomics preserves the spatial context of gene expression enabling us to study distinct tumor areas and the influence on overall diversity.

Our spatial transcriptomics analysis uncovered tumor clusters with distinct genetic profiles that showcase various functional areas in depth and offer valuable understandings into the diversity of ccRCC types. Some of these tumor clusters exhibited activity in genes responsible for protein catabolism and reduced abundance of genes related to mitochondrial respiration processes. We also show isoform expression within tumoroids, in particular glutaminase (GLS) especially with the prevalence of the highly metabolically active GAC isoform that is expressed in regions where mitochondrial gene abundance is lower; whereas the KGA isoform displayed a more focal expression pattern. Combining long-read spatial transcriptomics with organoid models presents a novel strategy for unravelling gene and transcript level complexity.

## 1. Introduction

Renal cell carcinoma (RCC) is a major health burden worldwide, with 434,419 new cases reported globally in 2022, resulting in 155,702 deaths ^1^. Clear cell renal cell carcinoma (ccRCC) is the predominant pathological subtype of renal cell carcinoma, accounting for approximately 70% of all RCC cases.^2^. Drug response in RCC is highly variable due to extensive intertumoral heterogeneity, making it challenging to predict which patients will benefit from specific treatments ^3,4^. While targeted therapies have been developed and approved in recent years, the effectiveness of these drugs remains inconsistent among individuals, highlighting the need for better strategies to identify actionable targets related to tumor heterogeneity and its microenvironment ^5^.

The development of precision medicine for RCC has been hindered by the lack of reliable preclinical models that accurately represent the genetic and structural features of patient tumors, with existing models such as cell lines and patient-derived xenografts (PDXs) having significant limitations ^6^. Organoid technology has emerged as a promising alternative, providing 3D models that closely mimic tumor structure and function; there has been a recent study that assessed mutational landscape, global gene expression profile, and cellular heterogeneity ^7^. However, it is worth noting that current tumor organoid (tumoroid) models often lack tumor microenvironment, including stromal and immune components, and do not incorporate spatial gene expression analysis.

Understanding the complexity of gene expression at the transcript level is crucial, as alternative splicing increases transcriptome complexity and plays essential roles in tumor development and progression ^8,9^. Spatial transcriptomics technologies, including the 10x Genomics Visium platform, have enabled high-throughput spatial mapping of gene expression in tumor tissues by capturing poly-adenylated RNA on spatially barcoded slides. However, these methods predominantly rely on short-read sequencing, which capture 3’ tags and are then subsequently digested to smaller fragments, thus missing critical information about full-length transcript variation.

Here, we describe a tumoroid culture system using adult ccRCC tissues that closely recapitulates the histological features of the parental tumors. Using this system, we successfully grew tumoroids and applied them to 10x Genomics Visium slides for spatial transcriptomics analysis. Additionally, we prepared barcoded libraries using the latest Oxford Nanopore Technologies Chemistry v14, enabling long-read sequencing to capture gene and isoform expression within the tumoroids with higher accuracy.

## 2. Materials and Method

### 2.1 Preparation of tumoroid samples

#### Materials and Reagents

Mayer’s Haematoxylin was purchased from pfm medical UK, Eosin Y 1% alcoholic and xylene were obtained from Cell Path.

##### Tumour tissue

clear cell Renal Cell Carcinoma (ccRCC) biopsies were obtained from partial or complete nephrectomy undertaken as curative treatment of kidney cancer. Ethical approval was granted by Lothian Biorepository (SR1787 10/ES/0061). Biopies were kept in DMEM / F12 Basal Medium which is made up by Advanced DMEM/F-12 (Gibco) supplemented with 10 mM HEPES (Gibco), 2 mM Glutamax (Gibco) and 1% Antibiotic antimycotic (Sigma) at 4°C before processing.

##### Tumoroids

Tumor tissue was cut into small fragments. One portion was fixed in 10% formalin for H&E and the remainder was digested in DMEM / F12 Basal Medium supplemented with 5mg/ml Collagenase type II (Gibco) and 10uM ROCK inhibitor (Tocris) at 37°C for 30 to 60 minutes on an orbital shaker. Digested tissue then went through a 70um cell strainer to obtain a suspension of single cells. Red Red Blood Cell (RBC) Lysis Buffer (Invitrogen) was added after to eliminate the RBCs. Then cells were washed three times by DMEM / F12 Basal Medium and a cell count was performed. Approximately 5-10 million cells were seeded into a 100 mL spinner flask containing 45 mL DMEM / F12 Basal Medium, 5 mL Fetal Calf Serum (10%), 10uM ROCK inhibitor, and 50ng/ml hEGF (Gibco). Spinner flask rotation speed was set at 27.5rpm and incubated at 37°C with 5% CO2.

#### Tissue processing, Embedding and Sectioning

For multiplex immunofluorescence (mIF) and H&E only, **p**rimary tissue was fixed in 10% Formalin overnight at room temperature. Tumoroids washed in PBS and fixed in 4% PFA 30 min at RT, then washed in PBS and were embedded in 2% melted agarose (100-200 uL). 70% EtOH is added to agarose, enough to cover the pellet (∼5mL) and then pellet is dislodged into the alcohol. Both tumoroid agarose pellets and tissue are processed overnight and embedded in paraffin.

### 2.2 Image Acquisition

After methanol fixation and H&E staining, the Visium slide was scanned at 20x magnification using the ZEISS Axioscan 7 with a custom scanning profile that includes tissue focusing and the capture of fiducial marks. The acquired images were saved in .czi file format. Each slide section was then exported as a high-resolution bigTIFF file for downstream analysis.

### 2.3 10x genomics Visium experiments

The Visium Spatial Tissue Optimization Slide & Reagent Kit (10x Genomics, Pleasanton, CA, USA) was used to optimize permeabilization conditions for OCT-embedded ccRCC tumoroids, including determining the optimal cycle number for the amplification step, which is critical for achieving optimal library yield. Following this, spatially barcoded full-length cDNA was generated using the Visium Spatial Gene Expression Slide & Reagent Kit (10x Genomics) according to the manufacturer’s instructions

### 2.4 Long-read nanopore sequencing

After amplification, 10 ng of barcoded spatial cDNA transcripts were used for nanopore library construction following the manufacturer’s protocol with some modifications. Briefly, the 10x Genomics Visium amplicons were tagged with biotin using the primers: [Btn]Fwd_3580_partial_read1 (5’-/5Biosg/CAGCACTTGCCTGTCGCTCTATCTTCCTACACGACGCTCTTCCGATCT-3’) and Rev_PR_partial_TSO_defined (5’-CAGCTTTCTGTTGGTGCTGATATTGCAAGCAGTGGTATCAACGCAGAG-3’). After annealing and amplification of the biotinylated oligonucleotides, the transcripts were captured using M280 streptavidin beads (Invitrogen, 11205D). The captured transcripts were further annealed and amplified using the cPRM primers from the SQK-PCS114 kit (Oxford Nanopore Technologies, ONT), followed by ligation of the Nanopore adapters. The prepared library was then loaded onto R10.4.1 PromethION flow cells and sequenced using the P2 Solo sequencer.

### 2.5 Long-read nanopore data processing

Nanopore libraries were sequenced using MinKnow software with default settings, and raw reads were saved in .pod5 file format. Basecalling of the raw reads was performed using the dorado super accurate (SUP) model (v0.7.0) on four Nvidia A100 GPUs, with parameters set to --minscore 7, --barcode-both-ends, and --no-trim. The unaligned .bam files generated by dorado were used as input for the epi2me-labs/wf-single-cell (v2.2.0) pipeline, utilizing the 2024 10x Genomics reference genome and the parameter --kit ‘visium:v1’. This workflow extracts cell barcodes and UMIs from 10x Genomics libraries, producing genome aligned tagged .bam files and generating gene and transcript count matrices for downstream analysis.

### 2.6 Spatial analysis

In order to obtain the 10x Genomics Visium spatial barcode positions, a custom BAM-to-FASTQ script, [bam2fastq.sh](https://github.com/MustafaElshani/Visium_ONT_analysis/blob/main/bam2fastq_ONT.sh), was generated. Briefly, the script used the tagged.bam file containing demultiplexed reads from epi2me-labs/wf-single-cell and generated FASTQ files compatible with 10x Space Ranger. The spaceranger count function was then used to analyze the FASTQ files, from which we obtained the tissue_positions.csv file. Raw gene and transcript expression matrices generated by epi2me-labs/wf-single-cell were subsequently processed using the Giotto Suite package (version 4.1.0) [Ref].

Within Giotto, spots were filtered based on defined thresholds, including a minimum expression threshold of 1.5, feature detection in at least 5 spots, and a minimum of 760 detected features per spot. A scree plot was then used to assess the number of principal components to retain for dimensionality reduction. Leiden clustering was performed with a resolution of 1.0, using the doLeidenCluster function, and the clustering was based on a shared nearest neighbor (sNN) network. Plots visualizing the results were generated using Giotto’s plotting functions. Gene and transcript-level differentials were identified using the Gini coefficient for spatial markers and the Scran method ^10^ for differential expression analysis.

### 2.7 Differential isoform detection

To identify transcript-level differences between Leiden clusters, a Giotto object was created using the raw transcript expression matrix, and the findMarkers_one_vs_all function with the “scran” method was used within Giotto. The identified markers were filtered to exclude single-transcript genes. Furthermore, the expression levels of transcripts were aggregated per gene to calculate isoform proportions within each cluster. Isoform switching was assessed by calculating the proportion difference of each isoform across clusters, and genes with substantial isoform proportion changes between clusters were identified as candidates for differential isoform usage.

## 3. Results

To demonstrate the feasibility of utilizing spatial transcriptomics and long-read sequencing in an in-house developed ccRCC tumoroid model, we used tumoroids that have been shown to maintain key histological features and elements of the tumor microenvironment, including immune cell markers. We successfully embedded nine tumoroids onto 10x Genomics Visium slides, followed by on-slide reverse transcription and PCR amplification and image capture, yielding sufficient barcoded transcripts for the construction of a Nanopore Chemistry v14. From this, we obtained approximately 105 million raw reads, with an average transcript N50 length of 750 bases.

### 3.1 Spatial Transcriptomics and Differential Expression Analysis of ccRCC Tumoroids

After basecalling, the wf-single-cell pipeline was used to demultiplex the reads and generate gene and transcript-level matrices, incorporating spatial barcode information. Leiden clustering of the spatial transcriptomics data revealed five major spatial domains across the nine ccRCC tumoroids, indicating the inter- and intra-tumoroid heterogeneity, despite all tumoroids being derived from the same patient.

Cluster 1 was primarily characterized by the elevated abundance of ribosome-associated genes, such as RPL41 and RPS7, which suggest active translation processes. This was further supported by GO Biological Process enrichment, which identified significantly enriched pathways related to translation and peptide biosynthesis (Figure 1C, Cluster 1).

**Figure 1.**
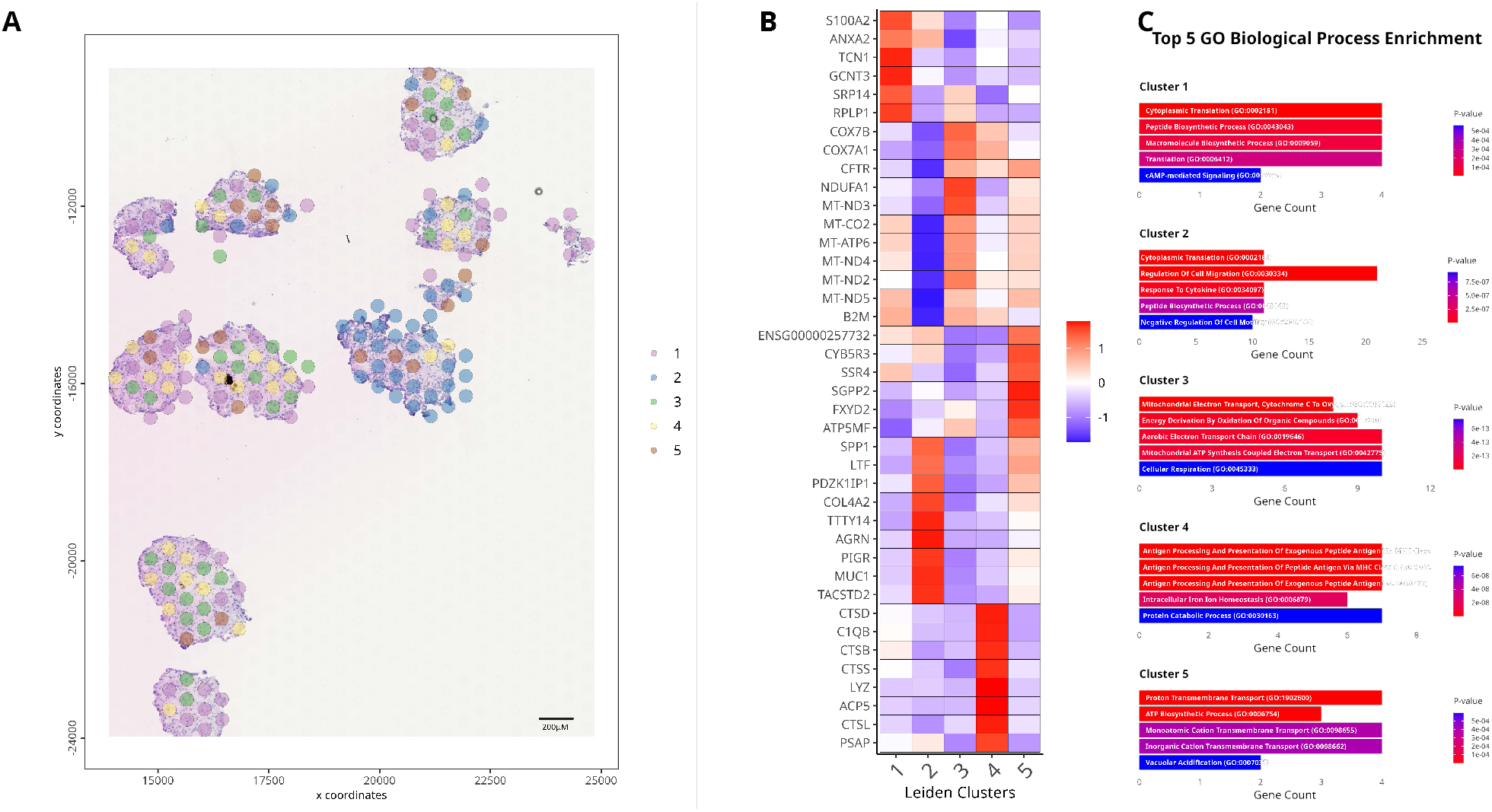
(A) Spatial distribution of 10x Genomics Visium spots overlaid on the H&E-stained image of tumoroids. Each spot represents spatial transcriptomic data, and the color-coding indicates distinct Leiden clusters identified through unsupervised clustering, highlighting regions with shared transcriptional profiles. (B) Heatmap showing the top 8 differentially expressed genes across Leiden clusters. The heatmap displays the scaled expression values, with red indicating high abundance and blue indicating low abundance within each cluster. (C) GO Biological Process Pathway Enrichment results generated using enrichR for each Leiden cluster. The top 5 pathways for each cluster are displayed based on their p-values, with red bars representing the pathways with the lowest p-values and blue bars representing the pathways with higher p-values. The gene counts in each pathway are shown along the x-axis.

Cluster 2 exhibited a distinct, higher abundance of genes related to cell migration, such as LTF, which aligns with histological observation of higher cellular density. The GO enrichment analysis further confirmed the involvement of cell migration and cytokine response pathways in this cluster (Figure 1C, Cluster 2). Notably, genes involved in the mitochondrial electron transport chain, including MT-ND3 and NDUFA1, were lower abundance in Cluster 2, contrasting with Cluster 3, where these genes were highly abundant (Figure 1B). This suggests potential metabolic differences between the clusters, with Cluster 3 displaying increased mitochondrial activity, which may indicate higher energy demand or oxidative metabolism (Figure 1C, Cluster 3).

Cluster 4, represented by yellow spots, displayed a higher abundance of genes involved in protein catabolism and antigen processing, including CTSB, CTSS, and CTSD (Figure 1B). These findings suggest an active role in proteolysis and immune response within this cluster. GO enrichment analysis highlighted the antigen processing and protein degradation pathways (Figure 1C, Cluster 4), and histological examination indicated that this cluster was predominantly localized in the interior regions of the tumoroids, hinting at a specialized microenvironment related to antigen presentation and intracellular protein catabolism.

Finally, Cluster 5 showed high levels of genes such as ATP5MF and ALDOA, which are associated with transmembrane transport and ATP biosynthesis (Figure 1B). This indicates that Cluster 5 is metabolically active, with enriched pathways related to energy production and transmembrane transport, likely supporting rapid cellular growth or stress response mechanisms (Figure 1C, Cluster 5).

Figure 2 shows the spatial distribution of the top differentially expressed genes as identified by the scran method, across the tumoroid sections. Mitochondrial genes, including MT-ND4, MT-ND3, MT-ATP6, MT-ND2, and COX7B, display distinct spatial expression patterns, with lower abundance predominantly observed in Cluster 2 and slightly elevated abundance in other tumoroids. This downregulation in Cluster 1 aligns with the earlier observation of reduced mitochondrial activity in this region, further confirming metabolic variability across the tumoroids. In addition, TACSTD2 and TTTY14 exhibited localized expression patterns, with TACSTD2 being expressed predominantly in one tumoroid.

**Figure 2.**
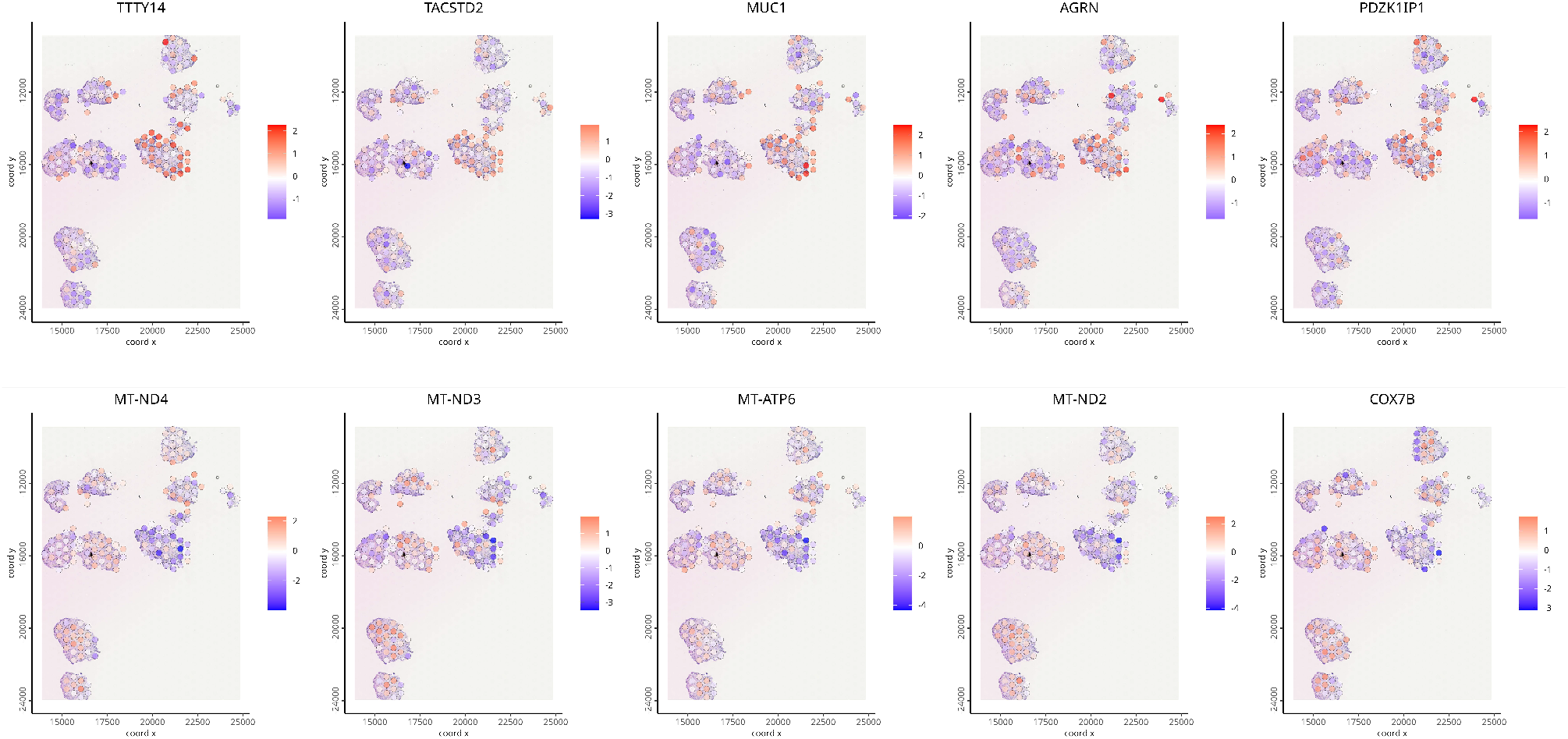
Spatial representation of the expression patterns of top differentially expressed genes as identified by the scran method. Each panel represents the spatial distribution of gene expression for a specific gene across the tumoroid sections. The color gradient, from blue to red, indicates the relative expression levels, with red representing higher expression and blue indicating lower abundance. The genes displayed include TTTY14, TACSTD2, MUC1, AGRN, PDZK1IP1, MT-ND4, MT-ND3, MT-ATP6, MT-ND2, and COX7B, illustrating varying expression levels across spatial clusters.

### 3.2 Gene and isoform expression in ccRCC tumoroids

The spatial expression patterns of the GAC isoform (Figure 3B), which encodes a more metabolically active form of the enzyme, show that it is primarily expressed in regions of the tumoroids with higher cellular density and histological activity. This suggests that the GAC isoform may be linked to areas of increased metabolic demand within the tumoroid microenvironment. In contrast, other tumoroids display significantly lower expression of the GAC isoform, indicating metabolic heterogeneity across the sample.

**Figure 3.**
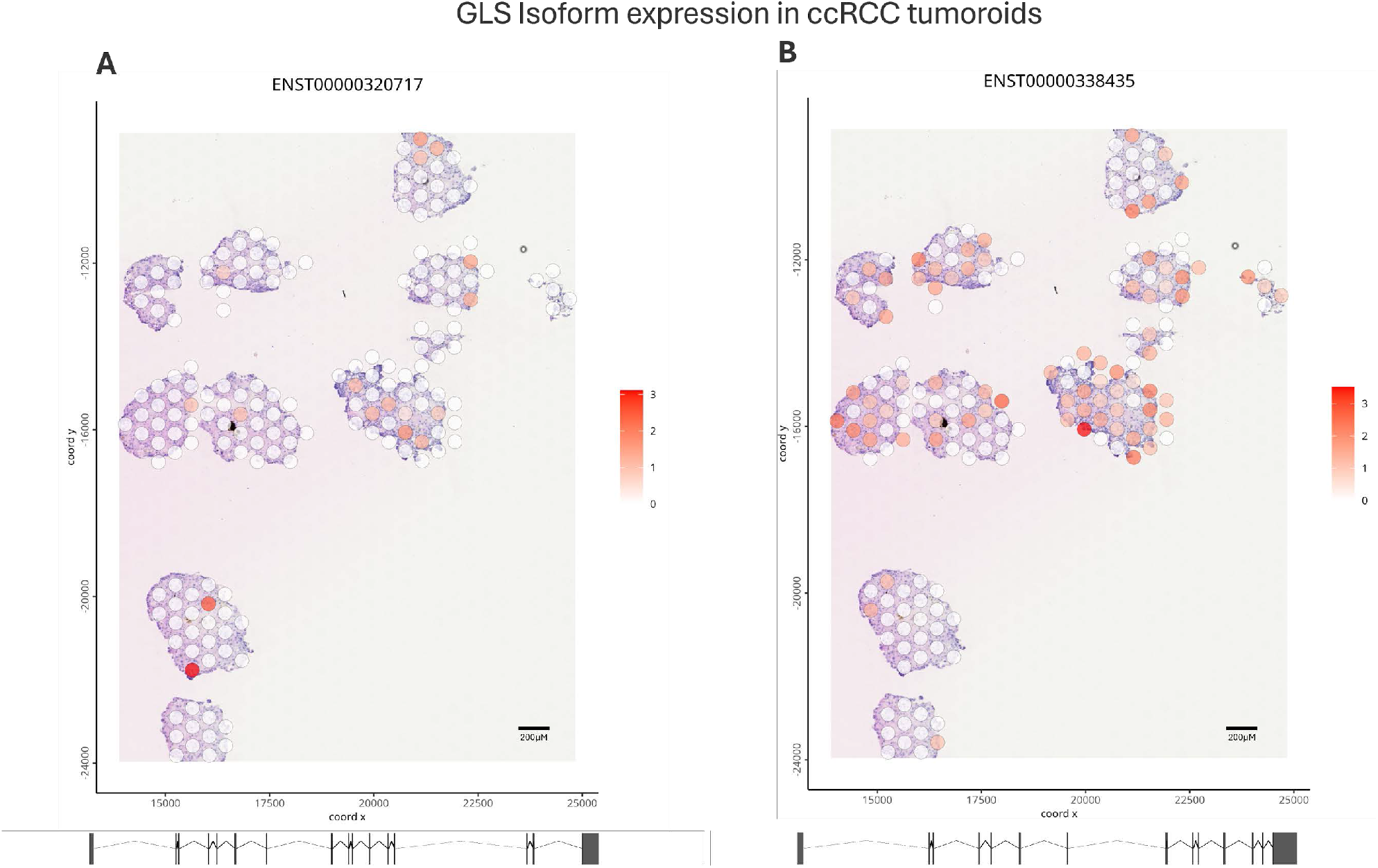
Spatial expression of GLS isoforms across ccRCC tumoroids. (A) Spatial distribution of the GLS KGA isoform (ENST00000320717), showing normalized expression across tumoroid sections. (B) Spatial distribution of the GLS GAC isoform (ENST00000338435), also showing normalized expression levels. The color scale, ranging from white (low expression) to red (high expression), indicates relative transcript abundance across the spatial coordinates of the tumoroids.

The KGA isoform (Figure 3A), although present at lower overall levels, shows focal expression in specific areas of the tumoroids. This localized expression suggests that the KGA isoform may play a specialized role in particular regions, potentially contributing to different metabolic states or microenvironmental conditions within the tumor.

These expression patterns highlight distinct spatial localization for each isoform within the ccRCC microenvironment, with varying levels of expression between regions. This spatial heterogeneity in isoform usage underscores the complex metabolic landscape of ccRCC tumoroids, suggesting that different regions within the tumor may rely on distinct glutaminase isoforms to meet their metabolic needs.

In the analysis of spatial isoform expression in ccRCC tumoroids, no significant isoform switching was observed between the defined Leiden clusters. However, a pronounced isoform heterogeneity was identified in specific regions of the tumor, with only a few spots showing high expression of particular isoforms. This suggests that certain isoforms within a gene are predominantly expressed in localized areas, indicating spatially restricted isoform usage by cellular subtypes.

These expression patterns highlight distinct spatial localization for each isoform within the ccRCC microenvironment, with varying levels of expression between regions.

## 4. Discussion

Tumor heterogeneity poses a significant challenge in understanding and treating ccRCC ^11^. Despite the increase in treatment options, patients’ responses are difficult to predict ^12,13^. Tumoroid models have shown promising avenues for recreating tumor heterogeneity^14,15^ in addition to being used for pre-clinical validation and drug efficiently^16,17^. Spatial transcriptomics profiling allows to map gene expression within its tissue setting while maintaining tissue architecture, tumor structure and specific patterns of diversity, in localized areas ^18,19^.

Our unsupervised clustering revealed multiple transcriptomic profiles, highlighting intra- and inter-tumoroid heterogeneity. It emphasizes the significance of transcriptomic profiling in dealing with the complexity of tumor heterogeneity. Notably, our findings reveal distinct transcriptional heterogeneity within the tumoroids and across tumoroids, which includes abundance of genes involved in pathway such as proteolysis and mitochondrial function.

Clear cell RCC is marked by significant metabolic reprogramming ^20,21^, and the development and survival of clear cell renal cell carcinoma (ccRCC) is significantly influenced by glutamine metabolism^22^. Initially, different tumoroids established from the same patient exhibit distinct transcriptomic profiles, highlighting intra-patient heterogeneity.

We demonstrate the usefulness of transcriptomics, and a workflow that allows study of tumoroids, in exploring the diversity in gene and isoform expression across tumoroids. Through this method, we have been able to chart out localized patterns of expression which give us information on how various sections of a tumor might utilize or require different metabolic and functional routes. By combining transcriptomics with read sequencing techniques we gain a deeper insight into the intricate molecular makeup of ccRCC tissues. This integrative approach offers a promising avenue for future research focused on the tumor microenvironment, alternative splicing, and the development of precision oncology strategies

## 5. Conflict of Interest

The authors declare no conflicts of interest.

## 6. Author Contributions

ME and DJH: conception and design of research. ME, YZ and HA performed experiments and analysed data. ME and DJH interpreted the results. ME performed bioinformatic analysis and generated figures. ME drafted the manuscript. All authors edited and revised the final version of the manuscript.

## 7. Funding

This work was supported by NuCana plc (ME, DJH), University of St Andrews Sanctuary Scholarship (HA) and the KATY project which has received funding from the European Union’s Horizon 2020 research and innovation program under grant agreement No 101017453 (DJH).

## 8. Acknowledgments

The authors acknowledge Research Computing at the James Hutton Institute for providing computational resources and technical support for the “UK’s Crop Diversity Bioinformatics HPC” (BBSRC grants BB/S019669/1 and BB/X019683/1), use of which has contributed to the results reported within this paper.

## Notes

### Competing Interest Statement

The authors have declared no competing interest.

## References

1. Bray, F., Laversanne, M., Sung, H., Ferlay, J., Siegel, R.L., Soerjomataram, I., and Jemal, A. (2024). Global cancer statistics 2022: GLOBOCAN estimates of incidence and mortality worldwide for 36 cancers in 185 countries. CA: A Cancer Journal for Clinicians 74, 229–263. 10.3322/caac.21834.

2. Sánchez-Gastaldo, A., Kempf, E., Alba, A.G.del, and Duran, I. (2017). Systemic treatment of renal cell cancer: A comprehensive review. Cancer Treatment Reviews 60, 77–89. 10.1016/j.ctrv.2017.08.010.

3. Barata, P.C., and Rini, B.I. (2017). Treatment of renal cell carcinoma: Current status and future directions. CA: A Cancer Journal for Clinicians 67, 507–524. 10.3322/caac.21411.

4. Chen, Y.-W., Wang, L., Panian, J., Dhanji, S., Derweesh, I., Rose, B., Bagrodia, A., and McKay, R.R. (2023). Treatment Landscape of Renal Cell Carcinoma. Curr. Treat. Options in Oncol. 24, 1889–1916. 10.1007/s11864-023-01161-5.

5. Bi, K., He, M.X., Bakouny, Z., Kanodia, A., Napolitano, S., Wu, J., Grimaldi, G., Braun, D.A., Cuoco, M.S., Mayorga, A., et al. (2021). Tumor and immune reprogramming during immunotherapy in advanced renal cell carcinoma. Cancer Cell 39, 649-661.e5. 10.1016/j.ccell.2021.02.015.

6. Yuan, Z., Fan, X., Zhu, J.-J., Fu, T.-M., Wu, J., Xu, H., Zhang, N., An, Z., and Zheng, W.J. (2021). Presence of complete murine viral genome sequences in patient-derived xenografts. Nat Commun 12, 2031. 10.1038/s41467-021-22200-5.

7. Li, Z., Xu, H., Yu, L., Wang, J., Meng, Q., Mei, H., Cai, Z., Chen, W., and Huang, W. (2022). Patient-derived renal cell carcinoma organoids for personalized cancer therapy. Clinical and Translational Medicine 12, e970. 10.1002/ctm2.970.

8. Hu, W., Wu, Y., Shi, Q., Wu, J., Kong, D., Wu, X., He, X., Liu, T., and Li, S. (2022). Systematic characterization of cancer transcriptome at transcript resolution. Nat Commun 13, 6803. 10.1038/s41467-022-34568-z.

9. Kahles, A., Lehmann, K.-V., Toussaint, N.C., Hüser, M., Stark, S.G., Sachsenberg, T., Stegle, O., Kohlbacher, O., Sander, C., Rätsch, G., et al. (2018). Comprehensive Analysis of Alternative Splicing Across Tumors from 8,705 Patients. Cancer Cell 34, 211-224.e6. 10.1016/j.ccell.2018.07.001.

10. Lun, A.T.L., McCarthy, D.J., and Marioni, J.C. (2016). A step-by-step workflow for low-level analysis of single-cell RNA-seq data with Bioconductor. Preprint at F1000Research, 10.12688/f1000research.9501.2 10.12688/f1000research.9501.2.

11. Hu, J., Tan, P., Ishihara, M., Bayley, N.A., Schokrpur, S., Reynoso, J.G., Zhang, Y., Lim, R.J., Dumitras, C., Yang, L., et al. (2023). Tumor heterogeneity in VHL drives metastasis in clear cell renal cell carcinoma. Sig Transduct Target Ther 8, 1–16. 10.1038/s41392-023-01362-2.

12. Motzer, R.J., Tannir, N.M., McDermott, D.F., Frontera, O.A., Melichar, B., Choueiri, T.K., Plimack, E.R., Barthélémy, P., Porta, C., George, S., et al. (2018). Nivolumab plus Ipilimumab versus Sunitinib in Advanced Renal-Cell Carcinoma. New England Journal of Medicine 378, 1277–1290. 10.1056/NEJMoa1712126.

13. Li, X., Shong, K., Kim, W., Yuan, M., Yang, H., Sato, Y., Kume, H., Ogawa, S., Turkez, H., Shoaie, S., et al. (2022). Prediction of drug candidates for clear cell renal cell carcinoma using a systems biology-based drug repositioning approach. eBioMedicine 78. 10.1016/j.ebiom.2022.103963.

14. Esser, L.K., Branchi, V., Leonardelli, S., Pelusi, N., Simon, A.G., Klümper, N., Ellinger, J., Hauser, S., Gonzalez-Carmona, M.A., Ritter, M., et al. (2020). Cultivation of Clear Cell Renal Cell Carcinoma Patient-Derived Organoids in an Air-Liquid Interface System as a Tool for Studying Individualized Therapy. Front Oncol 10, 1775. 10.3389/fonc.2020.01775.

15. Atanasova, V.S., de Jesus Cardona, C., Hejret, V., Tiefenbacher, A., Mair, T., Tran, L., Pfneissl, J., Draganić, K., Binder, C., Kabiljo, J., et al. (2023). Mimicking Tumor Cell Heterogeneity of Colorectal Cancer in a Patient-derived Organoid-Fibroblast Model. Cellular and Molecular Gastroenterology and Hepatology 15, 1391–1419. 10.1016/j.jcmgh.2023.02.014.

16. Botti, G., Di Bonito, M., and Cantile, M. (2021). Organoid biobanks as a new tool for pre-clinical validation of candidate drug efficacy and safety. Int J Physiol Pathophysiol Pharmacol 13, 17–21.

17. Ooft, S.N., Weeber, F., Dijkstra, K.K., McLean, C.M., Kaing, S., van Werkhoven, E., Schipper, L., Hoes, L., Vis, D.J., van de Haar, J., et al. (2019). Patient-derived organoids can predict response to chemotherapy in metastatic colorectal cancer patients. Science Translational Medicine 11, eaay2574. 10.1126/scitranslmed.aay2574.

18. Denisenko, E., de Kock, L., Tan, A., Beasley, A.B., Beilin, M., Jones, M.E., Hou, R., Muirí, D.Ó., Bilic, S., Mohan, G.R.K.A., et al. (2024). Spatial transcriptomics reveals discrete tumour microenvironments and autocrine loops within ovarian cancer subclones. Nat Commun 15, 2860. 10.1038/s41467-024-47271-y.

19. Arora, R., Cao, C., Kumar, M., Sinha, S., Chanda, A., McNeil, R., Samuel, D., Arora, R.K., Matthews, T.W., Chandarana, S., et al. (2023). Spatial transcriptomics reveals distinct and conserved tumor core and edge architectures that predict survival and targeted therapy response. Nat Commun 14, 5029. 10.1038/s41467-023-40271-4.

20. Pandey, N., Lanke, V., and Vinod, P.K. (2020). Network-based metabolic characterization of renal cell carcinoma. Sci Rep 10, 5955. 10.1038/s41598-020-62853-8.

21. Chakraborty, S., Balan, M., Sabarwal, A., Choueiri, T.K., and Pal, S. (2021). Metabolic reprogramming in renal cancer: Events of a metabolic disease. Biochimica et Biophysica Acta (BBA) - Reviews on Cancer 1876, 188559. 10.1016/j.bbcan.2021.188559.

22. Wang, M. (2023). Targeting glutamine use in RCC. Nat Rev Nephrol 19, 151–151. 10.1038/s41581-023-00684-2.

